# Immuno-Phenotyping of High-Grade Glioma Infiltrating Immune Cells Reveals Grade Specific Differences in Cells of Myeloid Origin

**DOI:** 10.1101/2020.09.26.314542

**Authors:** Raksha A. Ganesh, Jayashree V. Raghavan, Pranali Sonpatki, Divya Naik, Priyanka Arunachalam, Darshat Shah, Akhila Lakshmikantha, Shibu Pillai, Komal Prasad Chandrachari, Kiran Mariswamappa, L. D. Sathyanarayana, Nameeta Shah, Siddharth Jhunjhunwala

**Affiliations:** Mazumdar Shaw Center for Translational Research; Centre for BioSystems Science and Engineering, Indian Institute of Science; Mazumdar Shaw Medical Center

**Keywords:** Glioblastoma, Immunosuppression, Neutrophils, CD163, CD86, CD63

## Abstract

Gliomas are heavily infiltrated with immune cells of myeloid origin. Past studies have shown that high-grade gliomas have a higher proportion of alternatively activated and suppressive myeloid cells when compared to low-grade gliomas, which correlate with poor prognosis. However, the differences in immune cell phenotypes within high-grade gliomas (between grade III and IV) are relatively less explored, and a correlation of phenotypic characteristics between immune cells in the blood and high grade tumors has not been performed. Additionally, myeloid cells of granulocytic origin present in gliomas remain poorly characterized. Herein, we address these questions through phenotypic characterizations of monocytes and neutrophils present in blood and tumors of individuals with glioblastoma (GBM, grade IV) or grade III gliomas. Our data show that CD163 expressing M2 monocytes are present in greater proportions in GBM tissue when compared to grade III glioma tissue. In addition, we observe that neutrophils are highly heterogeneous among individuals with glioma, and a greater proportion of granulocytic myeloid-derived suppressor cells are present in grade III gliomas when compared to GBM. Finally, we show that the expression levels of CD86 and CD63 showed a high correlation between blood and tumor, and suggest that these may be used as possible markers for prognosis.

## Introduction

High-grade gliomas, the most common form of brain tumors, are associated with poor prognosis^1^. The current standard of care has met with limited success, possibly due to the high adjacent tissue infiltrative capacity^2,3^, treatment resistant cells^4^, low-medium mutational burden^5^, and cellular heterogeneity^6,7^ of glioblastoma (GBM). Immunotherapy, such as the use of antibodies against programmed cell death protein 1 (PD)-1^8^ that focus on preventing immunosuppression by regulatory T cells, have shown limited success^3^, which may be due to the diversity of immunosuppressive cells present in the GBM microenvironment.

In addition to regulatory T cells, multiple reports have highlighted the increased presence of alternatively activated or suppressive cells of myeloid origin in GBMs^9–11^. For example, larger numbers of myeloid-derived suppressor cells (MDSCs) have been observed in the blood^12–15^, and these cells are also enriched in the GBM microenvironment^5,16–18^. An enrichment of suppressive monocyte/macrophages (M2 or M0 phenotype)^10,19–21^ has also been observed. Further, it has been suggested that the monocytes in the GBM microenvironment may be phenotypically and functionally different^22,23^, and the same is possibly true for neutrophils^24–26^ too. Determining the prevalence of such immuno-modulatory cells of myeloid origin and characterizing their phenotype, could have implications for the development of new therapies.

In this context, herein, we aimed to address a few unanswered questions with regard to myeloid cells in individuals with gliomas: 1) if there are differences in the abundance of suppressive myeloid cells between grade III gliomas and GBM. Most studies suggest that the frequencies of suppressive cells are increased in GBM when compared to lower grade gliomas (grade II), but comparisons between GBM and grade III do not show significant differences^10,17^. 2) the characterization of granulocytes in the tumor microenvironment. Wang et al.^25^ demonstrated that cells with neutrophil gene signatures were present in GBM, and Chai et. al.^26^ showed that individuals with GBM had a greater proportion of neutrophilic-MDSC in the blood when compared to heathy controls. However, detailed immuno-phenotyping of these neutrophil populations has not yet been reported. And 3) if there is a correlation in the phenotype of suppressive myeloid cells between blood and tumor tissue of the same individual, similar to regulatory T cells^27^. Identifying phenotypes of immune cells in the blood that are predictive of glioma severity might help in making decisions with regard to clinical treatment and management. To answer these questions, we evaluated the phenotype (surface protein expression) of myeloid cells obtained from tumor resections and blood of individuals with gliomas using multi-color flow cytometry.

## Methods

### Ethics statement

Both human glioma tissue and blood samples were collected from individuals following informed consent at the Mazumdar Shaw Medical Center (MSMC), and all procedures were conducted in compliance with the approved protocol of the Institutional Review Board (No: NHH/MEC-CL-EA-1-2018-536). The diagnosis was confirmed by histopathological examinations by pathologists at the MSMC.

### Sample collection and preparation of single cell suspensions

Freshly resected tumor tissue was transported to the laboratory on ice within one hour of surgery in cold Roswell Park Memorial Institute 1640 culture medium (RPMI, Gibco, USA) media containing 1% antibiotics (Pen-strep, Thermo Fisher Scientific, USA). Samples were washed three times with cold phosphate-buffered saline (PBS) with antibiotics and minced using a scalpel in a 60-mm petri dish. The tissue fragments were digested in 30ug/ml accutase (Gibco) in 5ml of RPMI for 10-15 minutes at 37°C and dissociated by pipetting with a 1 ml-pipette 2–3 times. Dissociation was stopped by adding 10ml of RPMI, and the cell suspension was passed through 70-μm cell strainers (BD Falcon, USA). The single-cell suspension was washed twice with cold PBS, centrifuged, and used for fixable live-dead staining, as described below. About 3ml peripheral venous blood drawn from consented individuals was spun down at 500g for 5 minutes and plasma was saved. The cell pellet was subjected to RBC lysis using ACK lysis buffer (0.15M Ammonium Chloride, 10mM Potassium Bicarbonate, 0.1mM EDTA) for 10 minutes at room temperature in 1:10 (blood:lysis buffer) ratio by volume. Lysis was quenched using PBS solution containing 4mM EDTA, and the solution was centrifuged at 400g for 10 minutes at 4 degree. Supernatant was discarded and the pelleted white blood cells were suspended in PBS.

Blood and tumor cell suspensions were labelled with fixable live-dead stain (0.3 μl dye/100 μl volume of 1 million cell suspension) for 20 minutes at room temperature. Staining was quenched using 2 ml PBS containing 1% bovine albumin and 4mM EDTA (flow cytometry buffer), and cells were centrifuged at 400g for 4 minutes at 4°C. Cell pellet was fixed using 2% paraformaldehyde (prepared in PBS) while subjecting the tube to pulse-vortex. After 30 minutes, cells were washed and suspended in the flow cytometry buffer. Cell suspensions not stained with fixable live-dead dye were also fixed as described above, and used as controls.

### Immuno-phenotyping using flow cytometry

Cells were stained with a panel of antibodies (all from BD Biosciences, USA) described in Supplementary Table 1. Antibodies were added to fixed cell suspensions (made up in flow cytometry buffer) as per manufacturer’s instructions. Samples were incubated at 4°C for 30 minutes, and washed once to remove unbound antibodies. After washing, the centrifuged pellet was suspended in 300 μl flow cytometry buffer and run through a flow cytometer (BD FACS Celesta, USA). Single color controls were prepared using compensation beads (BD Biosciences) to which appropriate antibodies were added. Fluorescence-minus-one (FMO) controls were prepared using live-dead stained cell suspensions by removing one antibody at a time, and replacing it with its isotype during the antibody staining step. Flow cytometry data was analyzed using FlowJo (FlowJo LLC, USA). A minimum of 20,000 CD45^+^ live events were collected from each tumor and blood sample. A minimum threshold of 100 events was used to report percentage positive and MFI values. Single-cell and tSNE analysis of flow cytometry data are described in supplementary methods.

### Immunohistochemistry (IHC)

IHC was performed on formalin-fixed paraffin-embedded 3-micron tissue sections for selected proteins. Deparaffinization was carried out in xylene solution followed by rehydration with a series of graded alcohol, and antigen retrieval in Tris-EDTA buffer, pH 9.0. Endogenous tissue peroxidases were blocked with 3% hydrogen peroxide, and nonspecific binding was blocked with 10% bovine serum albumin (BSA) in PBS with 0.1% Triton X-100 (PBST, pH 7.6). The sections were then incubated for 2h at room temperature with primary antibodies - CD163 (Abcam, ab182422, 1:200), CD14 (Abcam, ab183322, 1:200), MPO (Invitrogen, PA5-16672, 1:500) followed by HRP-conjugated secondary antibody incubation for one hour and developed with 3, 3 - diaminobenzidine chromogen (DAB). Sections were counterstained with Mayer’s hematoxylin and mounted. IHC images were acquired and analyzed by a neuropathologist. IHC image analysis is described in supplementary methods.

### Statistical Analysis

Statistical analysis was performed using either Graphpad Prism version 5 or ‘R’. Student’s t test, one-ANOVA or two-way ANOVA, chi-square, or Wilcoxon tests were used for statistical comparisons.

## Results

### Clinical characteristics

Individuals undergoing craniotomy at the Mazumdar-Shaw Medical Center were recruited into this study, following informed consent. A total of 25 individuals (including three healthy volunteers from whom blood was drawn) were recruited (Table 1), of which samples from 17 individuals (six diagnosed with grade III IDH1 mutant glioma (Grade III), eight with GBM, and three healthy controls with no tumor) were used for further analysis. Clinical data of these individuals with GBM and grade III glioma (and one individual with benign tumor - meningioma) along with tumor staging is provided in Supplementary Table 2. Among the individuals with GBM, and grade III gliomas, an underrepresentation of females is observed, which may be attributed to the relatively small number of samples being analyzed in this study.

**Table 1:** Clinical characteristics of individuals recruited into this study. Tumor grade refers to the WHO glioma grade, and IDH refers to the isocitrate dehydrogenase mutation, both of which were determined by a pathologist.

|  | **Total**<br>**(25)** | **GBM (8)** | **Grade**<br>**III (7)** | **No**<br>**Tumor –**<br>**healthy**<br>**(3)** | **Grade II**<br>**and I (2)** | **Benign**<br>**(1)** | **Other**<br>**Tumors**<br>**(4)** |
| --- | --- | --- | --- | --- | --- | --- | --- |
| **Age**<br>**median**<br>**(10<sup>th</sup>, 90<sup>th</sup>**<br>**Percentile)** | 32 (23,<br>61) | 59 (19,<br>68) | 32 (29,<br>55) | 32 (29,<br>32) | 25 (24,<br>26) | 32 | 27.5 (22,<br>44) |
| **Sex** |  |  |  |  |  |  |  |
| **Female**<br>**(%)** | 7 (28%) | 2 (25%) | 0 (0%) | 1<br>(33.3%) | 0 (0%) | 1<br>(100%) | 3 (75%) |
| **Male**<br>**(%)** | 18 (72%) | 6 (75%) | 7<br>(100%) | 2<br>(66.6%) | 2<br>(100%) | 0 (0%) | 1 (25%) |
| **Tumor IDH**<br>**Status** | 18 | 8 | 7 | - | 2 | - | 1 (3 not<br>applicable) |
| **IDH1**<br>**Mutant (%)** | 6<br>(33.3%) | 0 (0%) | 6<br>(85.7%) | - | 0 (0%) | - | 0 (0%) |
| **Wild Type**<br>**(%)** | 12<br>(66.6%) | 8<br>(100%) | 1<br>(14.3%) | - | 2<br>(100%) | - | 1 (100%) |

### Frequencies of Immune Cells in Tumor and Blood

To profile the tumor microenvironment, a portion of the resected tumor was used for immunohistochemistry (IHC) based identification of cell types, and another portion was used to generate single-cell suspensions to be analyzed by flow cytometry. IHC was performed on five grade III and five GBM samples, to specifically identify myeloperoxidase (MPO) and CD14 expressing cells. MPO was used as a marker to identify granulocytes, and CD14 as a marker for monocytes. IHC based analysis shows an increased number of MPO (Figure 1A) and CD14 (Figure 1B) expressing cells in GBM compared to grade III gliomas, which potentially correlates with suggestions of increases in immune cell infiltration with increasing glioma grade^7^.

**Figure 1:**
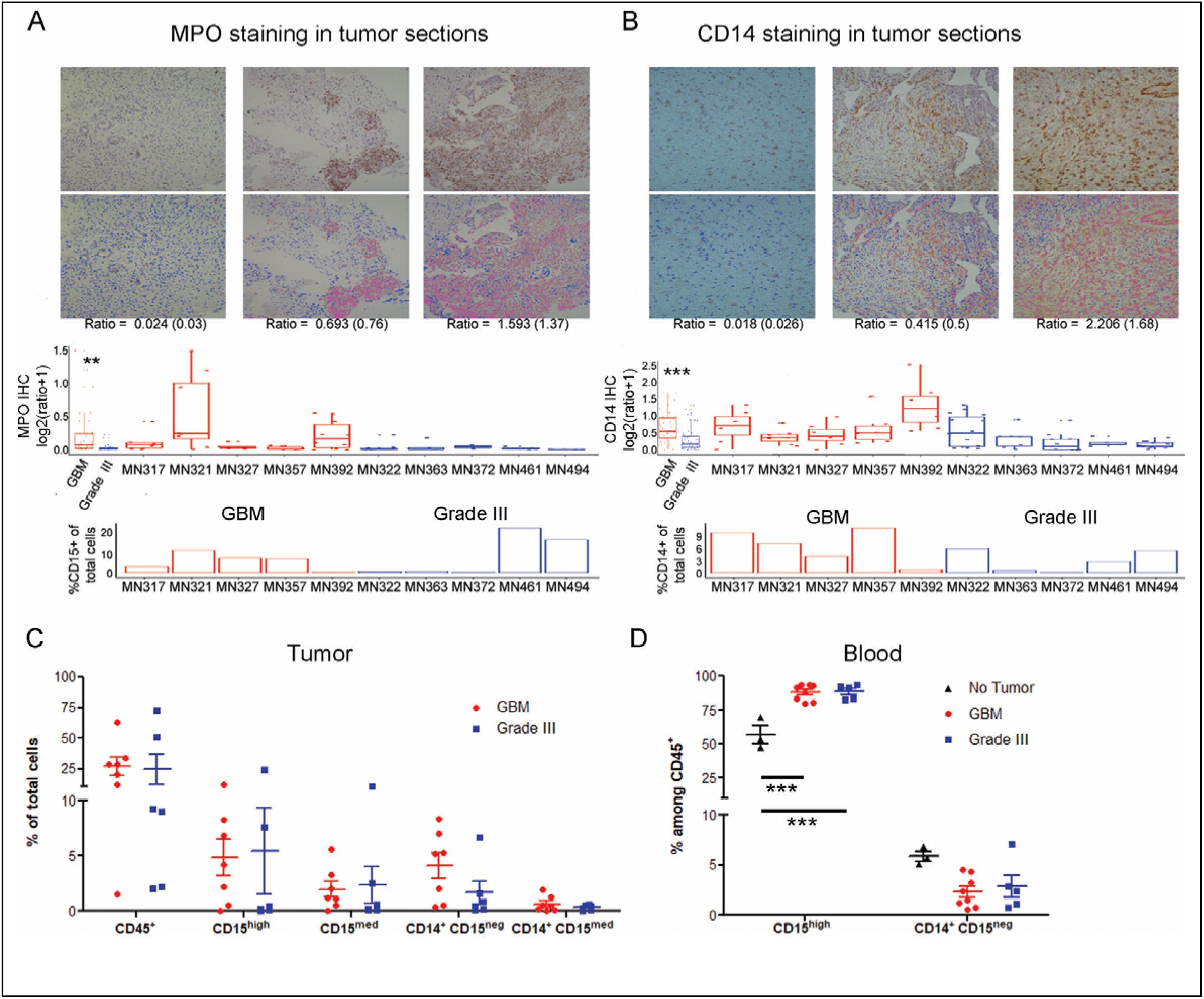
Immune Cell Frequencies. **A and B –** immunohistochemistry (IHC) based identification of myeloperoxidase (MPO) and CD14 expressing cells, respectively. A section from each tumor was stained for either MPO or CD14, images at 10x were taken from 5 different regions on each slide and the ratio of MPO/CD14 stain to the DAPI (nuclear stain) was determined using image analysis. Ratios (individual samples, and combined based on tumor grading) are shown in box plots. Images of three sections representing a low, medium and high ratio are shown as an inset in the graphs. Top panel of insets are images of IHC, and the bottom panel are digital conversions with pink representing the marker and blue nuclei for ease of viewing (enhanced contrast). In addition, percentages of CD15^+^ (neutrophils) and CD14^+^ (monocytes) cells determined by flow cytometry are shown below for comparison. **C –** percentages of immune cell subsets in tumors determined via flow cytometry. Percentages are calculated as the proportion of each subset among total live cells. Cell types were determined using the following markers: CD45^+^ – all immune cells; CD15^high^ – neutrophils; CD15^med^ – other granulocytes (gran.); and CD14^+^CD15^neg^ and CD14^+^CD15^med^ as two monocyte subsets. **D –** percentages of immune cell subsets in the blood. CD15^high^ – neutrophils and CD14^+^CD15^neg^ – monocytes. For statistical comparison of data, two-way ANOVA followed by Bonferroni’s test was performed. *** indicates p < 0.001. No significant difference was observed, if not indicated.

Separately, single-cell suspensions of resected tumors were obtained from six and seven individuals with grade III gliomas and GBM (tumor tissue was not available from one GBM individual for immuno-phenotyping), respectively, for flow cytometry based analysis. Single-cell suspensions were stained with antibodies that enabled the identification of total immune cells (determined as CD45 expressing live cells by flow cytometry), neutrophils (CD45^+^ CD15^high^), other granulocytes (CD45^+^ CD15^med^), and two different monocyte subsets (CD14^+^ CD15^neg^ and CD14^+^ CD15^med^) was quantified. Supplementary Figure 1A describes the gating strategies used to identify different immune subsets. Neutrophils and other granulocytes were distinguished by determining the expression of CD66b (present on neutrophils and absent on other granulocytes). Flow cytometry based analysis of percentages of immune cells revealed: (i) intra-tumor variability in immune infiltration when directly compared with histology sections (bottom panels of Figure 1A and 1B, where all CD15 and CD14 expressing cells are grouped together as granulocytes and monocytes, respectively); and (ii) inter-tumor variability in the frequencies of various immune cell subsets (Figure 1C). Nevertheless, the combination of histology and flow cytometry data are suggestive of an increased presence of myeloid cell subsets in the tumor microenvironment of GBM when compared to grade III gliomas.

In addition, peripheral venous blood was collected from three, five and eight individuals who were healthy (no tumor), had grade III gliomas, or GBM, respectively. In blood, unlike tumors, a single monocyte subset was observed (CD14^+^ CD15^neg^), and all the CD15 expressing cells were labeled as neutrophils, as cells expressing medium to low levels of CD15 were not observed (Supplementary Figure 2). Neutrophils were found to be present in significantly higher percentages in both GBM and grade III gliomas when compared to healthy controls (Figure 1D). This increase in neutrophil percentage resulted in a drop in monocyte percentages in the blood of individuals with tumors, but it was not significantly lower than that of healthy controls.

### Phenotyping Immune Cells in Tumor and Blood

To further characterize the myeloid cell subsets, single-cell suspensions of the tumor lysate and blood were analyzed via flow cytometry using four antibody panels that focused on neutrophil and monocyte/macrophage activation markers, as well as markers for MDSC (Supplementary Table 1). This analysis focused on determining the percentage of cells expressing a specific protein (as percentage positive cells) or measuring the extent of protein expression on a group of cells to report as median fluorescence intensity (MFI). Percentage positive cells were measured when a multimodal distribution of protein expression level on cells was observed, like in the case of certain monocyte proteins (for example CD163). Among granulocytes and some surface proteins of monocytes, the expression level of proteins on cells was unimodal, and hence MFI was quantified as a measure for expression. Among the cells from tumors, most proteins that we measured (CD11b, CD15, CD33, CD54, CD62L, CD63, CD282, and HLA-DR) did not show statistically different expression levels across tumor grades. A few surface receptors did have significantly different expression, such as CD16 expressed at higher levels on grade III neutrophils, and CD284 expressed at higher levels on one of the monocyte subsets on grade III tumors than GBM (Supplementary Figure 3). Among cells from blood, neutrophils from individuals with tumors showed significantly lower expression of a few activation markers (CD11b, CD16, CD54, and CD63) and L-selectin (CD62L), when compared to healthy controls (Supplementary Figure 4). However, these differences were not significant between the two grades of tumor. Similarly, a few surface proteins (CD54, CD282, and HLA-DR) were expressed at lower levels on monocytes from the blood of individuals with tumors, when compared to healthy controls (Supplementary Figure 5).

Notably, one of the surface proteins, CD163, showed a distinctly different pattern of expression. CD163 expression levels on both subsets of monocytes, measured as percentage positive cells and MFI, were significantly higher in GBM when compared to grade III tumors (Figure 2A). These differences were observed in IHC sections too, which on quantification showed a higher ratio of CD163 expressing cells among all other cells in GBM when compared to grade III tumors (Figure 2B). Markedly, the expression level of CD163 on monocytes present in the blood was not significantly different among the two grades of tumor and healthy controls (Figure 2C). Further, expression levels were not significantly different in the granulocyte subsets in tumor and blood (Figure 2D), which may not be surprising as granulocytes are not known to express CD163.

**Figure 2:**
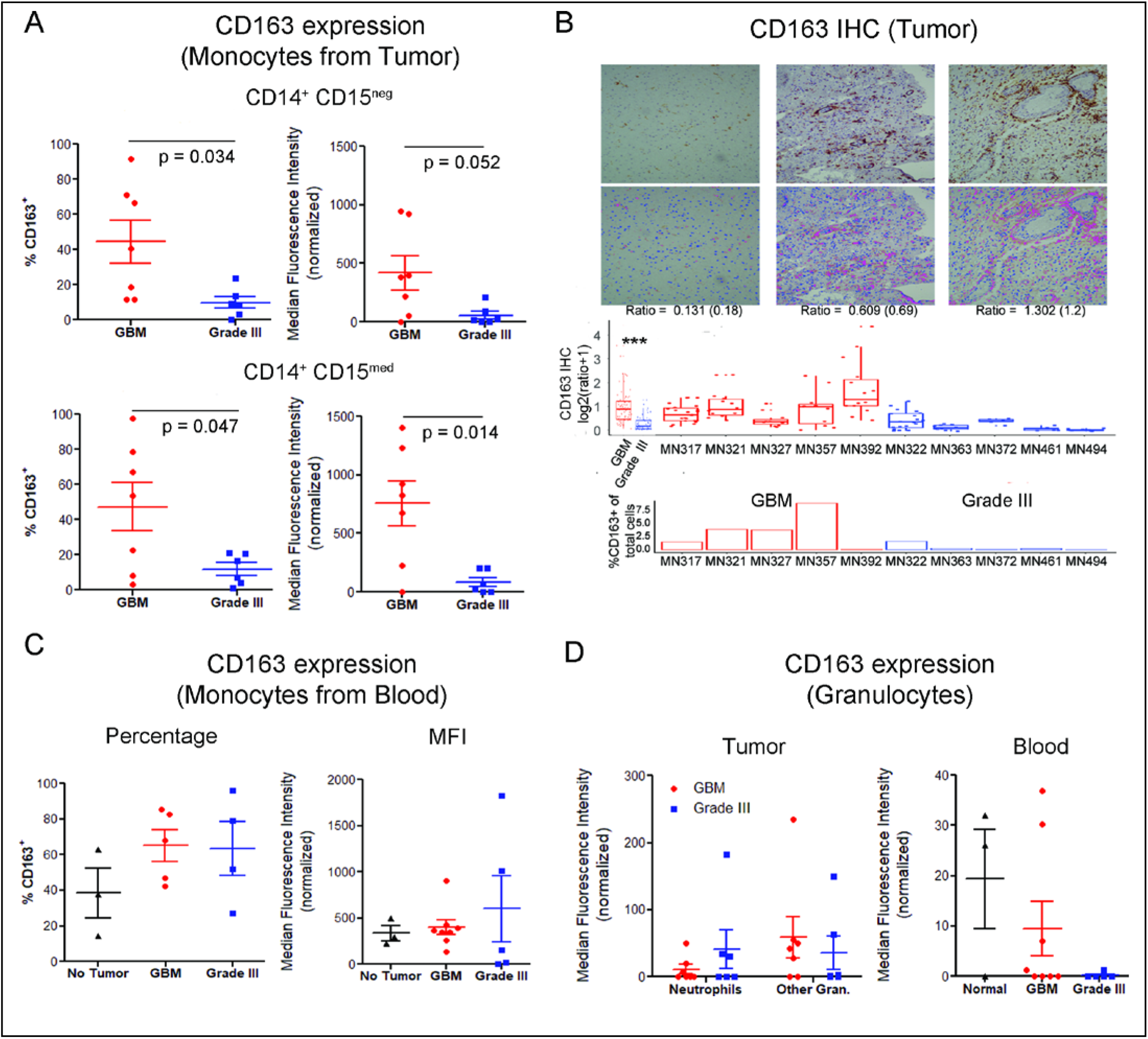
Expression of CD163. **A –** Expression levels among the two subsets of monocytes present in tumors. Left panel measures expression as a percentage of cells that are positive for CD163. Right panel measures expression as median fluorescence intensity (MFI) of the total subset population. For statistical comparison of data, two-tailed Student’s “t” test using Welch’s correction was performed. p values are indicated in the figure. **B –** Immunohistochemistry based determination of CD163 expression in tumor sections. Similar to other IHC images, a number of images from each section was obtained to determine the ratio of CD163 expressing cells among total cells (determined by counting nuclei). Ratios (individual, and combined based on tumor grading) are shown in box plots. Images of three sections representing a low, medium and high ratio are shown as an inset in the graphs. Top panel of insets are images of IHC, and the bottom panel are digital conversions with pink representing the marker and blue nuclei for ease of viewing (enhanced contrast). In addition, for the sake of comparison, the percentage of CD163^+^ cells as determined by flow cytometry for each tumor sample is also shown. *** indicates p<0.001 (ANOVA). **C –** Expression among monocytes in blood measured as either percentage positive or MFI. Significant differences were not observed between the 3 groups – one-way ANOVA. **D –** Expression of CD163 among neutrophils and other granulocytes (gran.) in tumor, and neutrophils in blood. Significant differences were not observed.

### Myeloid-Derived Suppressor Cells (MDSC)

MDSCs have been reported to be present in increased numbers in the blood^12–15^, and are enriched in the tumor microenvironment of individuals with GBM^5,16,17^. Utilizing the flow cytometry gating strategy described by Alban et. al.^17^, in a subset of our clinical samples (four individuals with GBM and grade III tumors and three healthy controls) MDSCs were identified as CD33 and CD11b expressing cells, but not expressing HLA-DR among the CD45^+^ immune cells (Supplementary Figure 6). We did not observe any significant differences in the frequencies of MDSC in the blood of individuals with tumors when compared to healthy controls (Figure 3A). Partitioning the MDSC into granulocytic-MDSC (CD15-expressing) and monocytic-MDSC (CD14-expressing) also did not reveal any differences.

**Figure 3:**
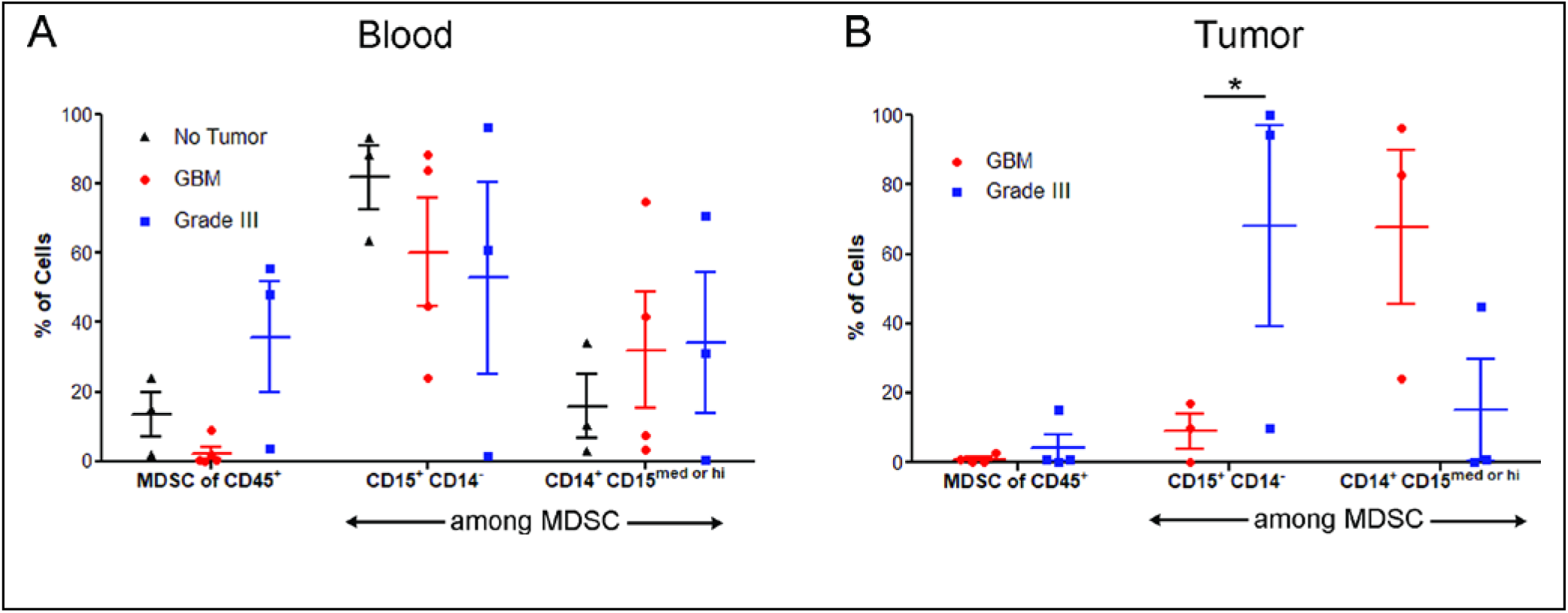
Myeloid Derived Suppressor Cell (MDSC) percentages in blood (A) and tumor (B). MDSC were determined as CD45+ HLA−DR− CD11b+ CD33+ cells (as shown in Suppl. Fig. 6). Granulocytic-MDSC were determined as CD15+ CD14− and monocytic-MDSC as CD14+ CD15med or high among the MDSCs. For statistical comparison of data, two-way ANOVA followed by Bonferroni’s test was performed. * indicates p <0.05, and where not indicated the differences are not significant.

Further, the overall MDSC frequency in the tumors was similar among the two grades of tumors. However, granulocytic-MDSC were enriched in grade III tumors (statistically significant), while monocytic-MDSC were enriched in GBM (not statistically significant) when compared to each other (Figure 3B). A point to note is that data on MDSC sub-populations (granulocytic and monocytic) were available for three of the four individuals (for tumors of both grades), as the number of events in one individual from each tumor grade was below our event threshold for analysis.

### Single-Cell Analysis

To reveal possible features in our data that traditional manual analysis may have missed, we performed a single-cell analysis of the immune cells (live CD45 expressing cells) using multi-flow cytometry single-event level data exported through FlowJo. We used the markers CD14 for monocytic lineage and CD15 for granulocytic lineage as well as the side scatter (SSC-A) as a measure of internal cellular complexity to classify cells into one of the twenty subsets (Figure 4A). For each of the parameters, we categorized the values as absent/low (−), medium (+), and high (++) based on their frequency distribution plots. Out of the 27 combinations we merged some to get the final twenty subsets (Supplementary table 3), of which we labeled eight based on current knowledge. We looked at the proportions of each of these 20 cell types among different groups in blood and in the tumor (Figure 4B). Similar to observations from the manual analysis, we found that compared to healthy controls, the blood of the individuals with gliomas (who were undergoing surgery) have a higher percentage of neutrophils and as a result fewer monocytes (Figure 4B). This can largely be attributed to the fact that the individuals undergoing surgery are on dexamethasone, which increases the proportion of neutrophils. In addition, we see that among the cell subsets SSC-A^−^CD14^−^CD15^+^, SSC-A^−^CD14^+^CD15^+^, SSC-A^−^CD14^+^CD15^−^ are higher in blood from healthy controls compared to blood from glioma patients, whereas NeutrophilsCD14^−^ cells are found in higher proportions in blood from individuals with glioma. We did not observe significant differences between blood from Grade III and GBM patients except for SSC-A^−^CD14^+^CD15^−^ cells, which are found in higher numbers in Grade III (Figure 4B and 4C).

**Figure 4:**
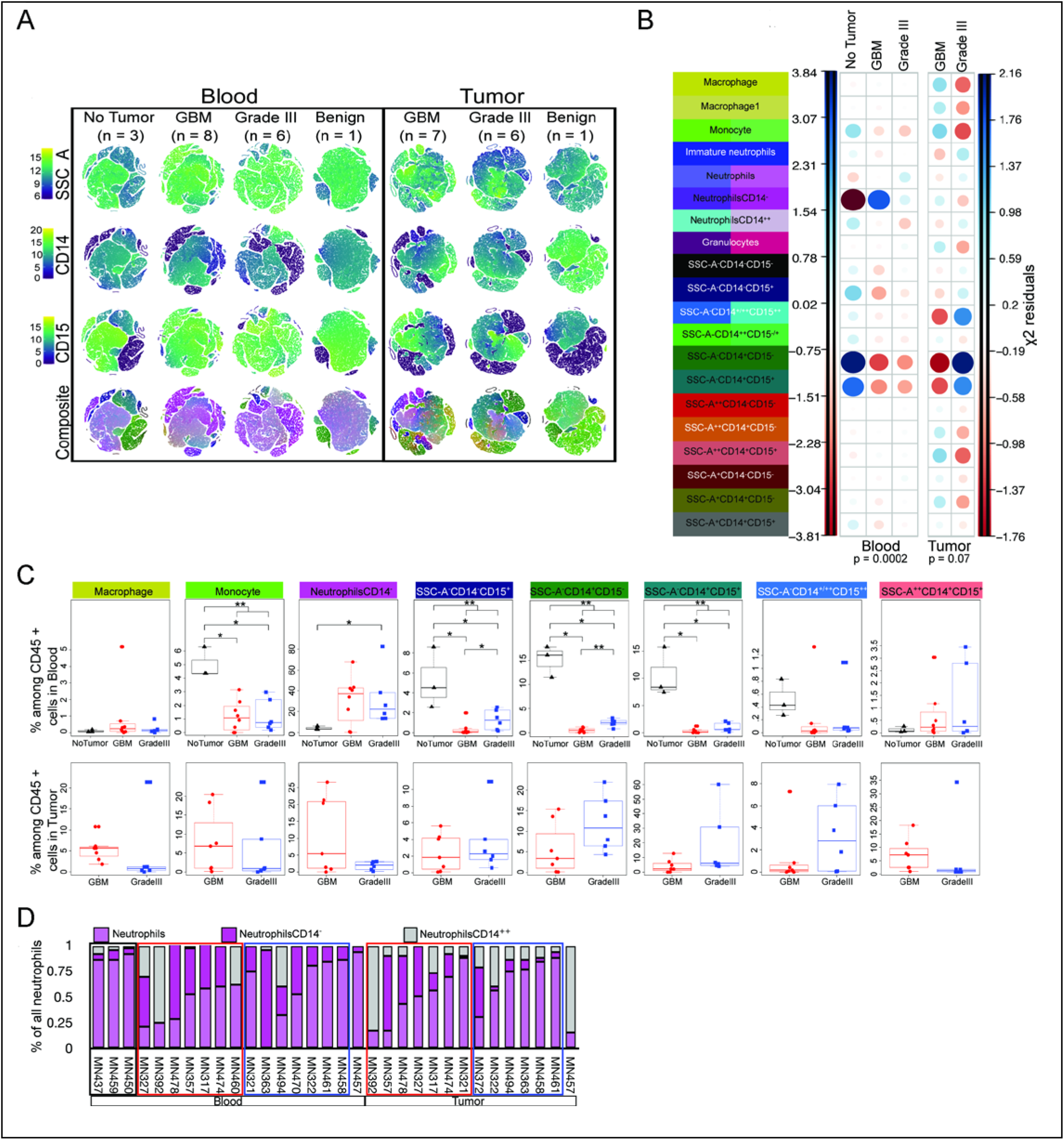
Single Cell Analysis. **A –** tSNE analysis was done across four groups in blood and three groups in tumor tissue. 50,000 events were chosen randomly and equally from all the samples in a group and tSNE was performed using SSC-A, CD14, and CD15 values. The composite figure was generated by merging the grey-scale images for SSC-A (Red), CD14 (Green), and CD15 (Blue) for visualization of different cell proportions among CD45+ live cells across groups. Immune cell composition is much more heterogeneous in tumor tissue compared to blood. The heterogeneity of immune cell types based on the three parameters of SSC-A, CD14, and CD15 is highest in the GBM tumor tissue as seen in the composite figure. **B –** Cell type proportions across blood and tumor tissue samples. Live CD45^+^ cells were classified into 20 cell types based on their SSC-A, CD14, and CD15 values and their proportions among CD45^+^ cells was calculated in each sample. For each cell type the median proportion was used for each of the groups. ***X***^2^ residuals were used to show the proportions of each subset in blood and tumor tissues, and the p values were calculated using a ***X***^2^ test. **C –** Boxplots of eight cell types showing different proportions across groups in blood and tumor tissue as seen in panel B. The Wilcoxon rank sum test was used to calculate p values, and * = p<0.05, ** = p<0.01. **D –** Percentage of three types of neutrophils; traditional neutrophils, neutrophilsCD14^++^, and neutrophilsCD14^−^ among all neutrophils is shown for all samples.

With regard to the immune cells infiltrated in GBM and grade III tumors, we observed that the cells in GBM tend to have higher SSC-A indicating higher granularity compared to grade III tumors (Figure 4A and 4B). We also observe a larger number of cell types in GBM compared to Grade III. We do not see statistically significant differences in different proportions, but we observe a trend for macrophages, monocytes, SSC-A^++^CD14^+^CD15^+^ to be higher in GBM tissue compared to Grade III, whereas SSC-A^−^CD14^+/++^CD15^++^, SSC-A^−^CD14^+^CD15^+^, SSC-A^−^CD14^+^CD15^−^ tend to be higher in Grade III (Figure 4B and 4C). One of our observations was the variability of CD14 expression on neutrophils (SSC-A^+/++^CD15^++^). CD14 expression displayed a trimodal distribution with no expression (CD14^−^), medium level expression (CD14^+^) and high expression (CD14^++^). The majority of neutrophils from healthy donors’ blood as well as from a patient with benign tumor (meningioma) expressed a medium level of CD14 expression (CD14^+^), which we have labeled as neutrophils. The percentage of NeutrophilsCD14^−^ and NeutrophilsCD14^++^ was significantly increased in the blood of glioma patients (Figure 4C and 4D). Also, these neutrophil subsets were found to be infiltrated in glioma tissue (Figure 4D). A subset of patients with GBM had a high percentage of NeutrophilsCD14^−^ (5/7) among all neutrophils in their tumor tissue as well as blood, in contrast to Grade III patients where we saw only one patient with a high percentage of NeutrophilsCD14^−^ (blood was not analyzed for this patient).

### Correlation of Blood and Tumor Immuno-phenotype

One possible outcome of such phenotypic analyses is the identification of markers in the blood that might help predict the grade of the tumor or the expected progression of the disease. To determine if such biomarkers are present, the phenotype of immune cells in the blood was compared to the phenotype of the same subset of immune cells present in the tumor, for each individual. Such a comparison was first performed through a correlation analysis of the expression levels (measured as MFIs) of various surface proteins among traditional monocytes (CD14^+^CD15^−^) and neutrophils (CD15^high^) Among monocytes, CD33, CD54, and CD86 showed a positive correlation, whereas, among neutrophils CD62L, CD63 and CD86 showed a positive correlation (Figure 5A). Performing correlation analysis among the cell-types identified by single-cell analysis also resulted in a high positive correlation for CD86 (Figure 5B and Supplementary Figure 7) across cell-types. High expression of these two markers is associated with poor prognosis based on both TCGA and CGGA data (Figure 5C). Additionally, we observed that the correlation of CD163 expression between blood and tumor was extremely poor, suggesting that their expression levels in the blood are unlikely to represent the expression levels in the tumor.

**Figure 5:**
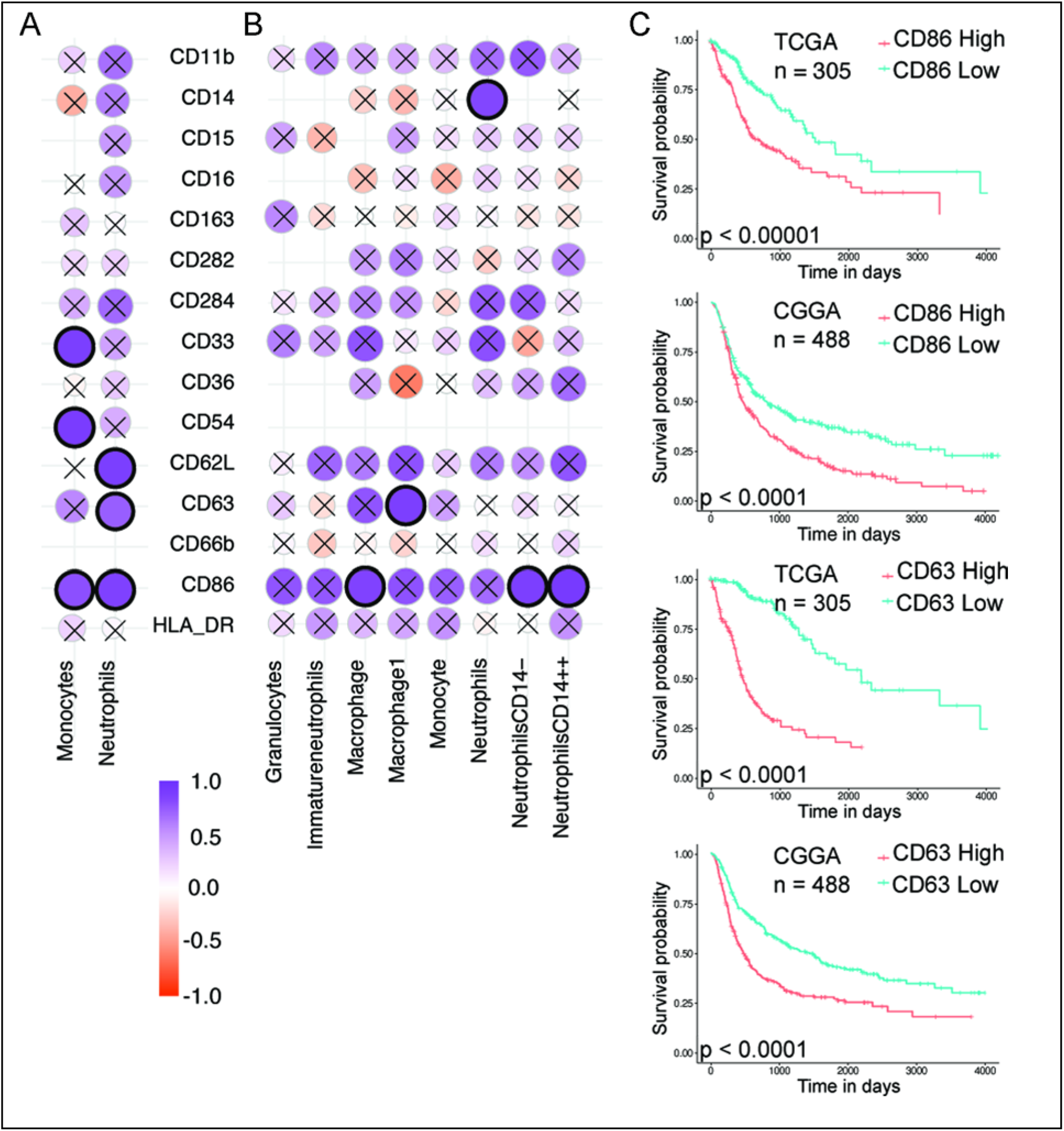
Correlation Analysis. **A –** Spearman’s correlation of surface protein expression levels (MFI) among neutrophils and monocytes in the blood and tumor. Correlation with p-value < 0.01 are highlighted in black circles. **B –** Spearman’s correlation between blood and tumor for eight cell types identified using single-cell analysis. **C –** Kaplan–Meier survival analysis for CD86 and CD63, markers showing correlation between blood and tumor values in both traditional flow cytometry analysis and single-cell analysis. Grade III with IDH mutation and Grade IV without IDH mutation samples from TCGA and CGGA data were used for this analysis. The samples were divided based on the median expression values of CD86 and CD63 genes.

A similar correlation analysis was performed on the frequencies of MDSC too. This analysis revealed that for both grades of tumors, the levels of monocytic-MDSC found in the blood to tumors are highly correlated (Supplementary table 4). Given the small number of individuals in each group, the implication of such correlations are modest at best.

## Discussion

There is a growing appreciation for myeloid cells in tumor microenvironments, especially those that are alternately activated or have suppressive functions. This is especially true for brain tumors, where cells of myeloid origin make up a large percentage of cells in the tumor^28–31^. Characterizing the number, phenotype and function of these myeloid cells has the potential to enhance current treatment strategies, or help develop new therapeutic approaches. Through the data presented here, we add to the current knowledge of myeloid cell phenotypes in high-grade gliomas.

First, we report that about one-fourth of the glioma cell count comprises of immune cells identified based on the expression of CD45, and about half of these cells could be classified as granulocytes or monocytes based on the expression of CD15 (or MPO in histology) and CD14, respectively. These numbers are lower than what has been reported historically^32,33^ (reviewed in^28,29^), which could be due to heterogeneity in the tumor or possibly due to population level variations. Additionally, in the glioma microenvironment, we observe an additional monocyte sub-population that we characterize as CD14^+^ CD15^med^ through manual analysis of flow cytometry data. Their overall proportions are rather low, and they phenotypically resemble the traditional monocytes (CD14^+^ CD15^neg^) present in the tumor microenvironment. This specific cell subset is not observed in the blood, and hence we speculate that they arise from traditional monocytes in the tumor microenvironment.

Phenotyping of the monocyte subsets in the tumor specifically revealed an enrichment of CD163 expressing cells in GBM compared to grade III gliomas. However, we did not observe any differences in the CD163 expression among the monocytes present in the blood of the same individuals. In fact, we did not observe a higher proportion of CD163 expressing monocytes in the blood of individuals with glioma when compared to healthy controls too, which appears to be contrary to previous reports by Heimberger and colleagues^19^ and Agrewala and colleagues^21^. But it is important to note the following differences: the former study compared the expression levels of CD163 at the RNA level, while the latter study used a different gating strategy (CD11b vs. CD14 used by us). Additionally, if we specifically look at the data related to the percentage expression of CD163 among monocytes in blood, we observe an increase (not statistically significant) in both tumor grades when compared to healthy controls. Importantly, we did not observe a correlation between CD163 expression levels in the blood and in the tumor of the same individuals, which might suggest that the tumor microenvironment plays an important role in either recruiting or converting activated monocyte/macrophage populations to CD163 cells (with possible suppressive function).

With regard to MDSC frequencies, we did not observe differences in the blood of individuals with tumors when compared to healthy individuals. This is in stark contrast to a number of previous reports that show increased numbers of these cells in the blood when compared to healthy controls^12–15,17,19^. One possible reason for this difference is that our study is underpowered for MDSC analysis, as data has been collected from a relatively low number of individuals (4 in the tumor groups and 3 in healthy controls). Another, and we speculate that the most likely possibility, is that our analysis was performed on whole blood, and not the PBMC fraction. How the use of whole blood instead of the PBMC fraction might impact the MDSC data is unclear, and further studies are necessary to determine the true reason for the differences. However, analysis of MDSC levels in the tumors reveals an interesting pattern of increased monocytic-MDSC in GBM and increased granulocytic-MDSC in grade III gliomas, when compared to each other.

Through the single cell analysis, we observe that neutrophils in both the tumor tissue and the blood of individuals with glioma are heterogeneous, as determined by their expression of various cell surface proteins. Neutrophil heterogeneity is now well-recognized^34^, although they remain poorly characterized. In the context of tumors, Singhal et. al^35^ had identified a unique sub-population of antigen-presenting cell like neutrophils in lung cancers. Our analysis also reveals a sub-population of neutrophils that express antigen-presenting cell receptors like CD86. In addition, we also observe heterogeneity among neutrophils (identified as CD15^high^ and CD66b^+^) based on CD14 expression levels, and specifically show that neutrophils that express high levels of CD14 (NeutrophilsCD14^++^) are enriched only in gliomas. The roles of these neutrophil sub-populations, and their effect on glioma progression remains to be studied.

Finally, by comparing the phenotype of myeloid cells between blood and tumor of the same individual, we were able to determine if blood phenotype is a representation of the tumor phenotype. To a large extent, CD86 and CD63 were the only cell-surface proteins whose expression levels correlated among myeloid cells in the blood and tumor. Given that higher expression of both these proteins in the tumor is associated with poor prognosis, we propose that the expression levels of these proteins on myeloid cells in the blood may be used as a prognostic marker for the progression of gliomas.

The following caveats apply to the data presented here. First, the tumor tissue was randomly divided into three parts, and the flow cytometry based phenotyping and immunohistochemistry analysis were performed on different parts. The data presented here may be influenced by the heterogeneity of gliomas. Second is that a majority of our results rely on immuno-phenotyping, and studies to understand cellular function have not been performed. The third is the absence of comparisons of suppressive immune cell presence with clinical outcomes. Correlation analysis of the same is planned for future studies.

## Acknowledgements

Anjali Vijaykumar and Shruti KS for helping in sample preparation for flow cytometry, and Yogesh Pasupathy and Anurag CN for sample collection.

