## Supplementary for "Immuno-Phenotyping of High-Grade Glioma Infiltrating Immune Cells Reveals Grade Specific Differences in Cells of Myeloid Origin"

**Table of Contents:**


| | |
|---|---|
| Supplementary Methods | 2 – 5 |
| Supplementary Figures | 6 – 14 |
| Supplementary Tables | 15 – 19 |

### Supplementary Methods

**Flow cytometry data export:** For single-cell analysis, compensated $CD45^+$ live cells were exported as FCS files group-wise for every patient using FlowJo. Single-cell data was extracted from FCS files using the Flowcore package in R and stored as CSV files for further analyses.

**tSNE analysis:** A pseudo count of 1 was added to all the intensity values and they were log2 transformed for downstream analysis. Events were randomly selected from samples to collectively form 50,000 events for each sample using the sample() function in R. These events were used to perform tSNE for each sample using the Rtsne() function with default parameters. Only the intensity values of SSC-A, CD14, and CD15 were used for tSNE calculations in individual samples. The log2 intensity values along with the tSNE dimension values were combined for each sample i.e. no tumor, GBM, Grade III, and Benign. ggplots package was used to visualize the tSNE data and the coloring of each event was based on the log2 intensity values of SSC-A, CD14, and CD15 where dark blue color indicated low expression and yellow color indicated high expression. A composite tSNE image was generated using ImageJ[1] by merging the grey-scale images for SSC-A (red), CD14 (green), and CD15 (blue) to visualize the representation of each cell type in blood and tumor tissue across groups.

**Cell type assignment and proportion calculation:** The cells were divided based on the SSC-A, CD14, and CD15 log2 intensities values into 20 subtypes. The proportions of each cell type in a sample was calculated as a percentage among CD45+ live cells. The benign subtype was excluded from the downstream analysis as it consisted of a single sample. The Wilcoxon rank-sum test in R was used to calculate the statistical significance of proportion differences across groups in blood as well as tumor tissues (Intensity_boxplots.pdf).

**Corrected Fluorescence minus one (FMO) calculation**

FMO values for a specific marker (for example, CD11b) was measured by determining MFI of this marker among the neutrophils, traditional monocytes, different monocytes, or granulocytes populations (based on gating) in tubes containing all antibodies except for the antibody against that marker (for example FMO for CD11b). The FMO values were subtracted from the intensity value to get the FMO corrected intensity. The antibodies for which FMO was not calculated were not included in this analysis. For blood and tissue samples, the FMO corrected intensities were calculated as follows:

- cell type = neutrophils/neutrophilsCD14-/neutrophilsCD14++

  FMO corrected intensity = intensity - FMOneutrophils

- cell type = macrophage/macrophage1

  FMO corrected intensity = intensity - FMOtraditional monocytes

- any other cell type

  FMO corrected intensity = intensity - max(FMOneutrophils, FMOtraditional monocytes)

- For tumor tissue samples the  FMO corrected intensities were calculated as follows:

  cell type = neutrophils/neutrophilsCD14-/neutrophilsCD14++

  FMO corrected intensity = intensity - FMOneutrophils

- cell type = macrophage

  FMO corrected intensity = intensity - FMOtraditional monocytes

- cell type = macrohage1

FMO corrected intensity = intensity - FMOdifferent monocytes

- cell type = granulocytes

  FMO corrected intensity - intensity - FMOgranulocytes

- any other cell type

  FMO corrected intensity = intensity - max(FMOneutrophils, FMOgranulocytes, FMOtraditional monocytes, FMOdifferent monocytes)

The FMO corrected intensities were not calculated for CD14 and CD15 antibodies. The boxplots were generated for each cell type and each protein across all samples (markerplots folder). The FMO corrected intensity values were then averaged for each cell type per sample. One-way ANOVA was used to find cell types with differential FMO corrected intensities across groups in blood and tumor tissue samples (markerplots folder).

**IHC image analysis:** IHC stained samples were digitized first by capturing the images at 10X resolution and viewed in OpenCV software. The goal of image processing was to separate the hematoxylin stained blue-purple hued nuclei from DAB stained brown cells and compute the ratio of respective areas occupied in the image. The images had differences in background color, so the focus was on identifying the foreground pixels for blue and brown stains. It was followed by finer post-processing of the identified regions to trim noise due to either a) lack of sharp boundary between foreground and background in some of the images or b) some background pixels also showing colors that we wanted to separate. Images were converted to HSV color space, and pixels belonging to blue hue (105-150 range chosen for blue) and brown hue (165-179, 0-15 ranges chosen for red) were identified. Occasionally depending on background variation, the hue ranges were adjusted slightly automatically with rules described next. These heuristics were derived from the visual examination of nuclei color in selected images. For brown stains, Value (V of HSV) of 127 was used as a cutoff to identify darker brown areas. In the case of blue stains, the region of interest (ROI) was converted to greyscale followed by contouring and thresholding to separate background noise from nuclei boundaries. Since nuclei were not of predictable shape (often wispy or amorphous), using shape-based identification was not an option, the procedure relied on thresholding heuristics to best match visual verification. Finally, masks were created for both stain types, and the absolute count of pixels belonging to either mask was used for computing the ratio of brown pixel count to blue pixel count. Notably, the intensity of blue or brown did not affect the ratio, only the number of pixels identified as belonging to the respective hue. (github location of the script – https://github.com/MSCTR/DigitalPathology/blob/master/process_bluebrownmasks-hsv.ipynb)

### Supplementary Figures

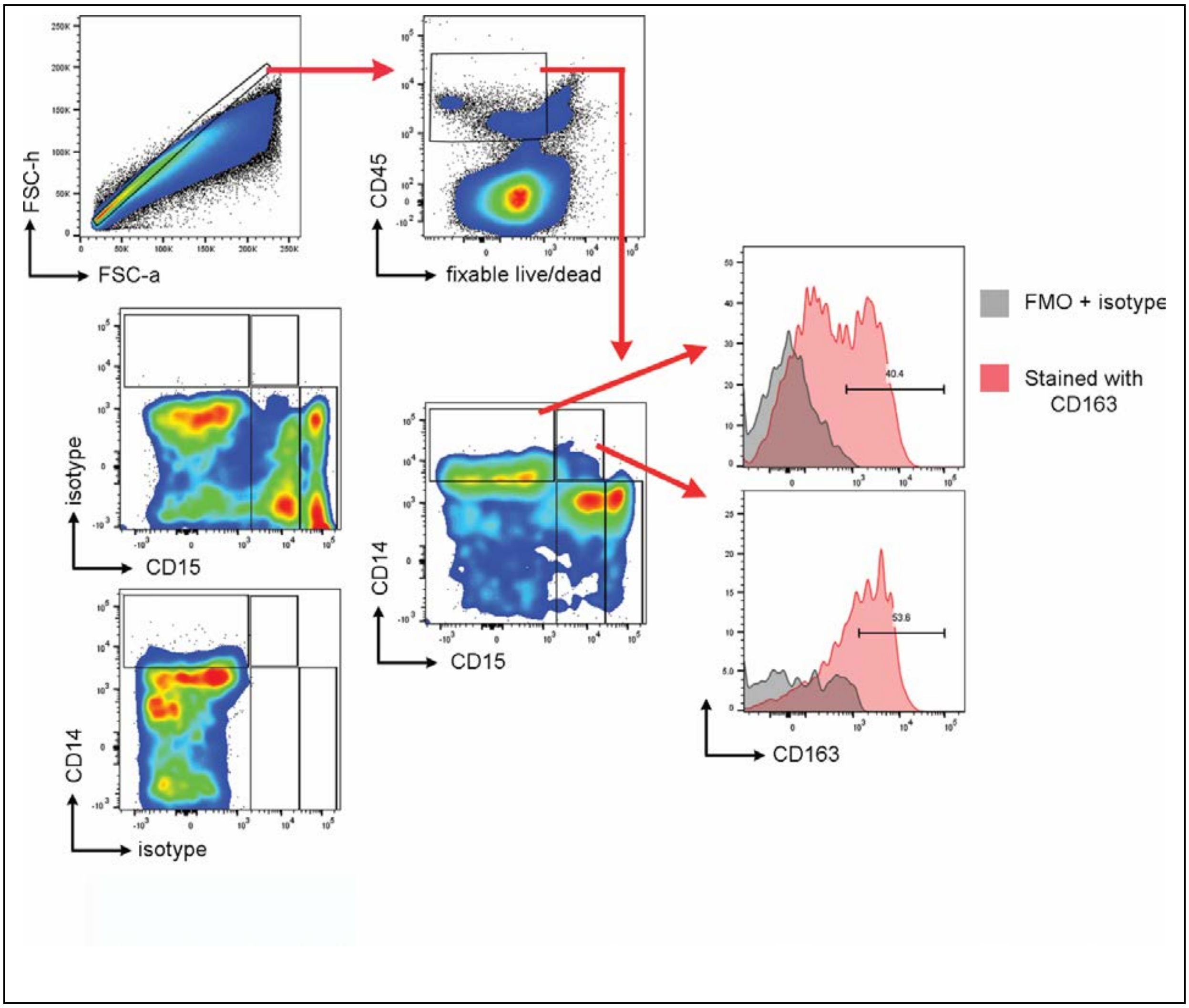


**Supplementary Figure 1:** Gating strategy used to analyze immune cells present in the tumor microenvironment. Prior to singlet gating (FSC-a vs. FSC-h), cells were gated based on forward and side scatter (not shown). Gates were drawn based on fluorescence minus one + isotype (labeled as isotype in this figure)

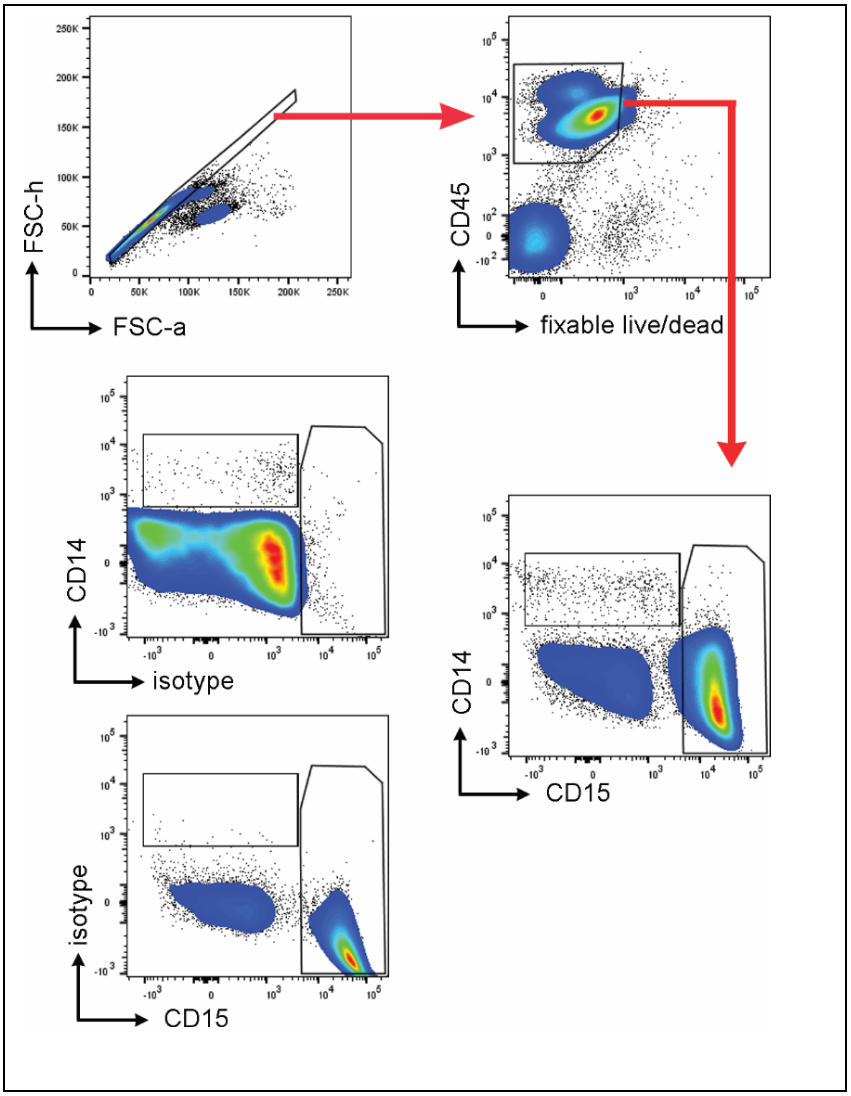


**Supplementary Figure 2**: Gating strategy used to identify specific immune cell subsets in the blood. Prior to singlet gating (FSC-a vs. FSC-h), cells were gated based on forward and side scatter (not shown). Gates were drawn based on fluorescence minus one + isotype (labeled as isotype in this figure)

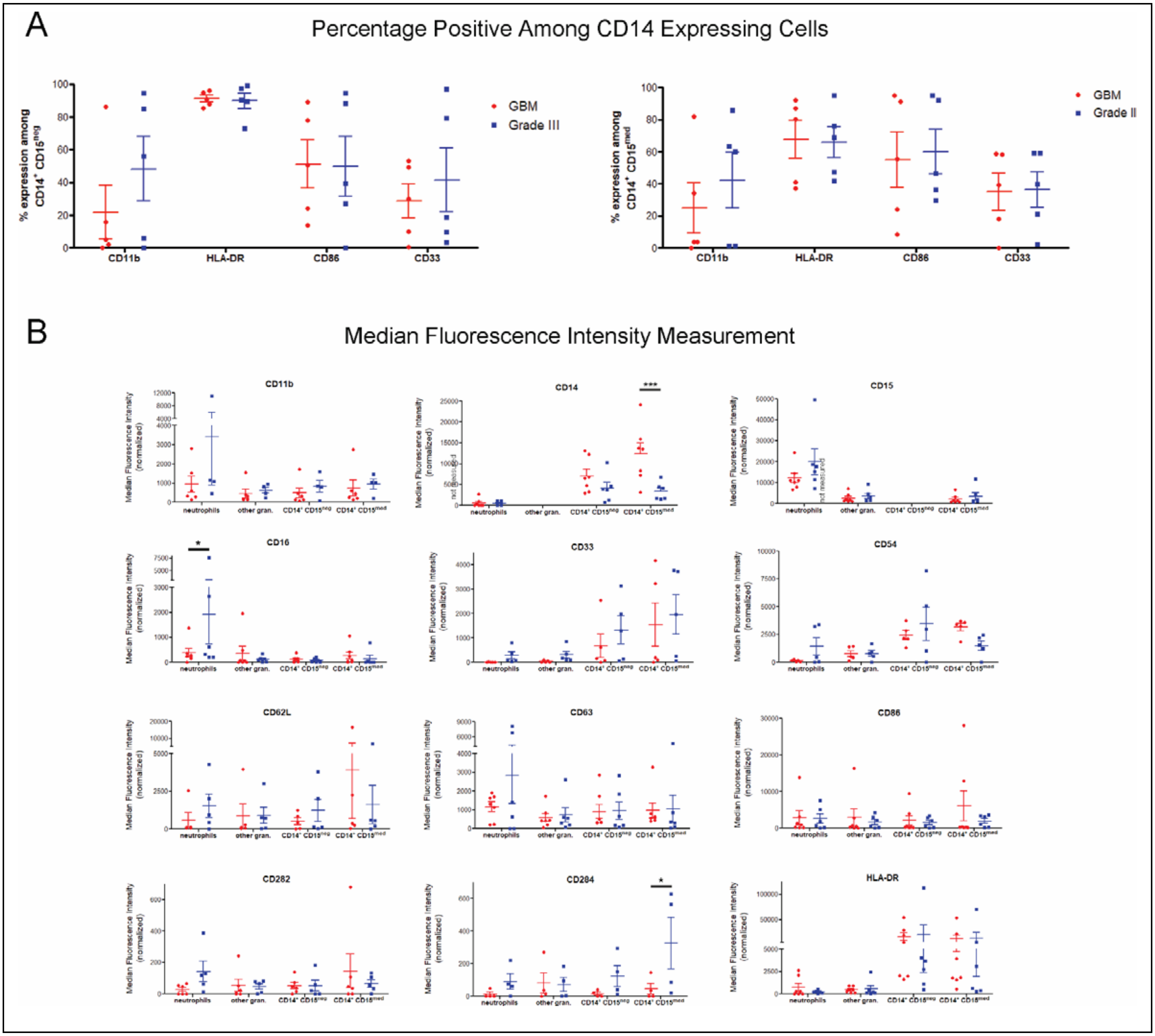


**Supplementary Figure 3:** Phenotyping of myeloid cell subsets present in tumors. Expression of a specific surface protein is presented as either: A – percentage positive cells; or B – median fluorescence intensity (MFI). Percentage positive cells were only determined for CD14 expressing cells, as they showed bimodal expression patterns for certain markers. Other cell subsets (neutrophils and other granulocytes) did not show bimodal expression of any markers. Hence only MFI was determined for these subsets. For statistical comparison of data, two-way

ANOVA followed by Bonferroni’s test was performed. * and *** indicates $p < 0.05$ and $p <0.001$, respectively. No significant difference was observed, if not indicated.

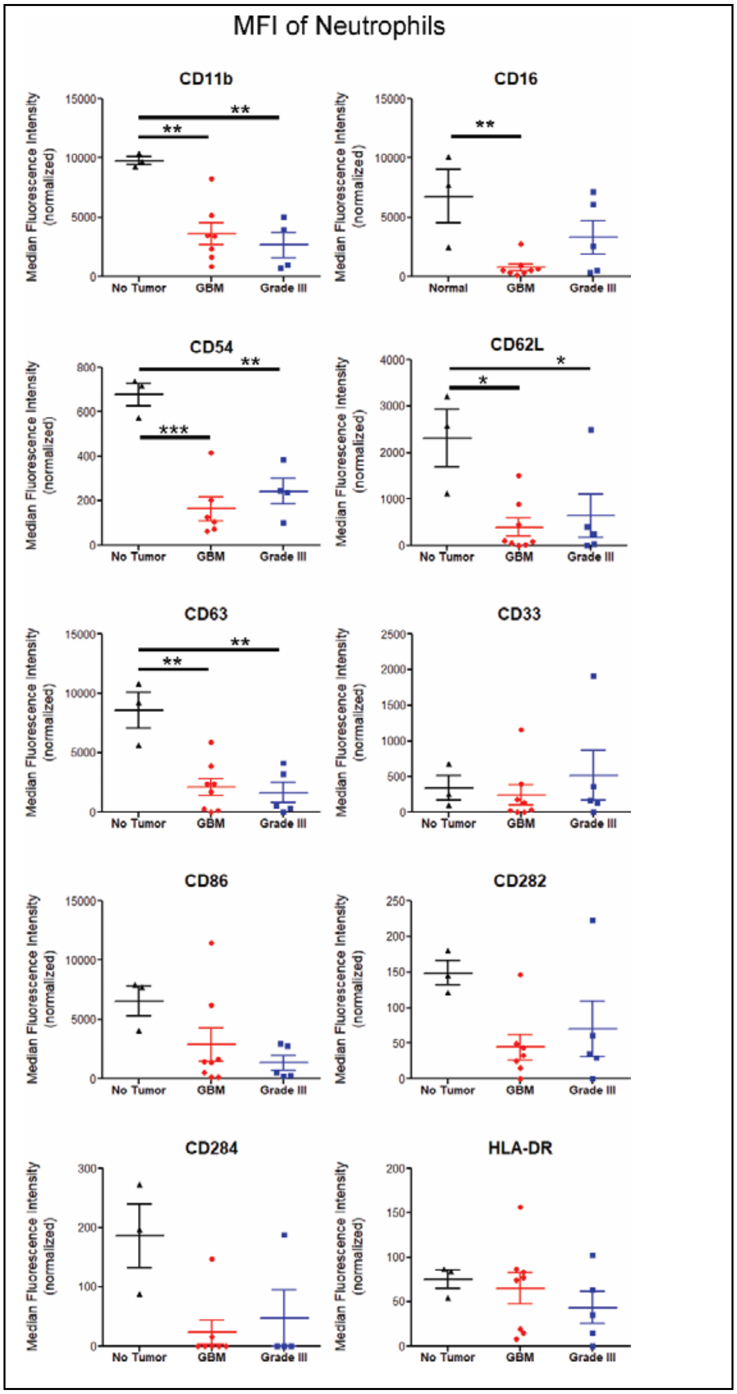


**Supplementary Figure 4:** Phenotyping of neutrophils in blood through median fluorescence intensity (MFI) measurements of specific surface proteins. For statistical comparison of data, one-way ANOVA followed by Tukey test was performed. *, **, and *** indicate $p < 0.05$, $p < 0.01$, and $p < 0.001$, respectively. No significant difference was observed, if not indicated.

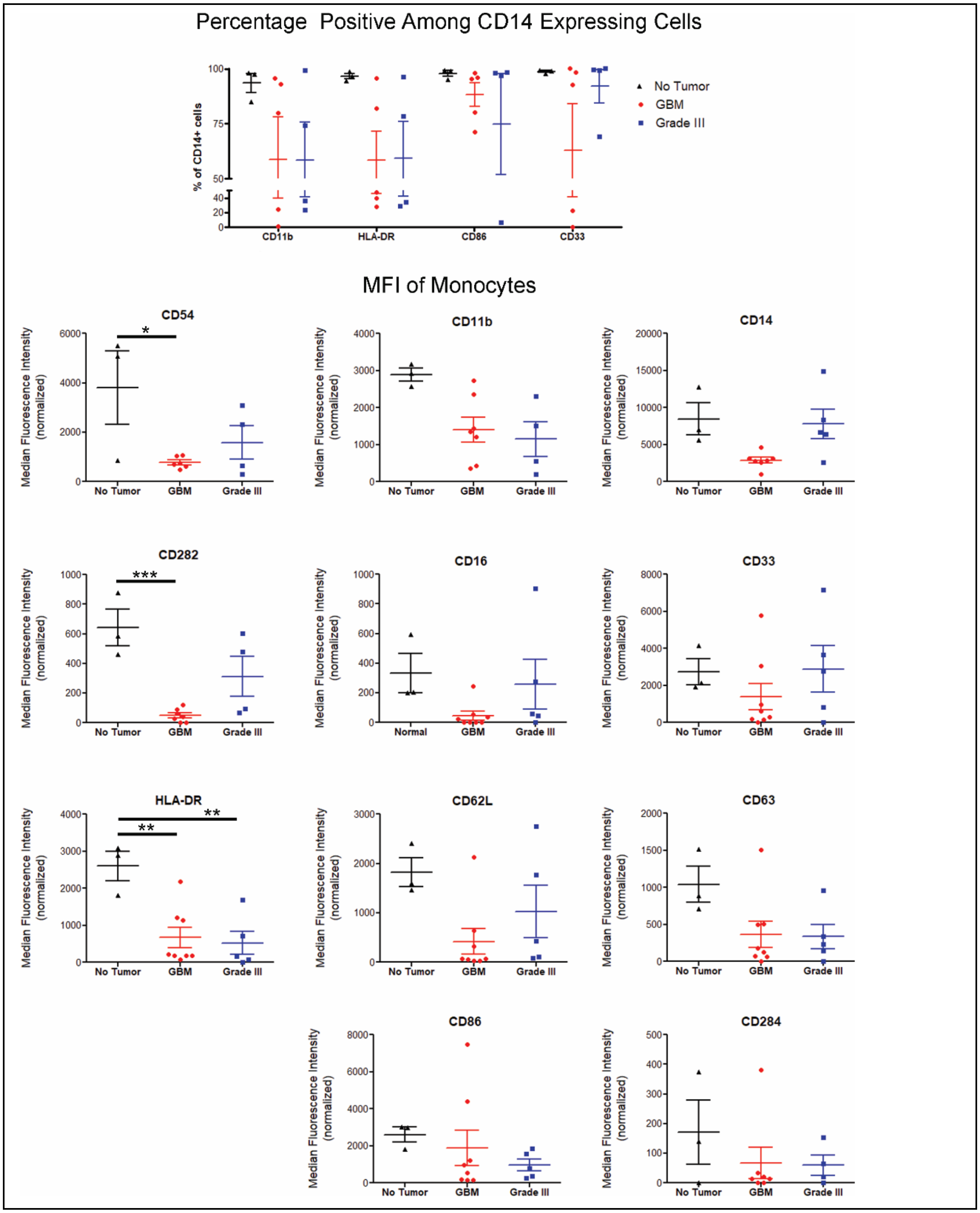


**Supplementary Figure 5:** Phenotyping of monocytes in blood through percentage or median fluorescence intensity (MFI) measurement of specific surface proteins. For statistical comparison of data, one-way ANOVA followed by Tukey test was performed. *, **, and ***

indicate $p < 0.05$, $p < 0.01$, and $p < 0.001$, respectively. No significant difference was observed, if not indicated.

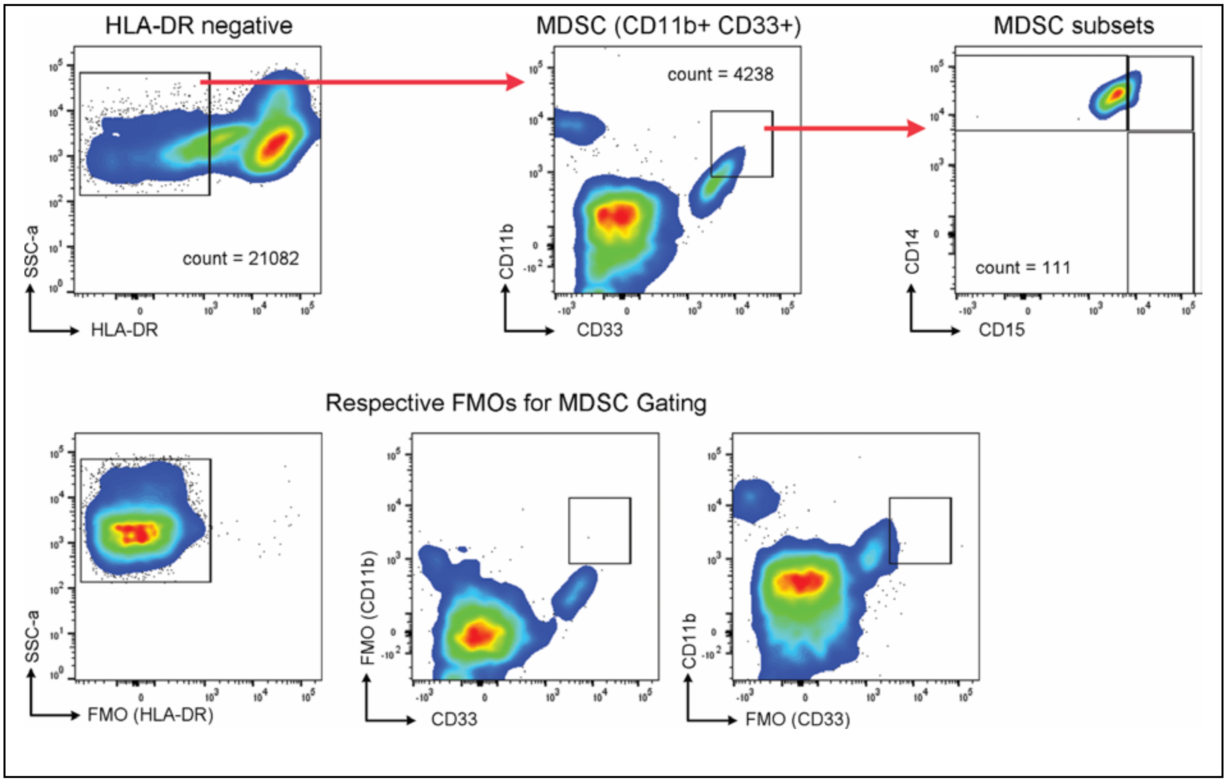


**Supplementary Figure 6**: Gating strategy used to identify myeloid derived suppressor cells (MDSC) and their subtypes. HLA-DR negative gating is on CD45+ live cells as shown in supplementary figure 1. Gates were drawn based on fluorescence minus one + isotype (labeled as FMO with the missing antibody in parentheses for this figure)

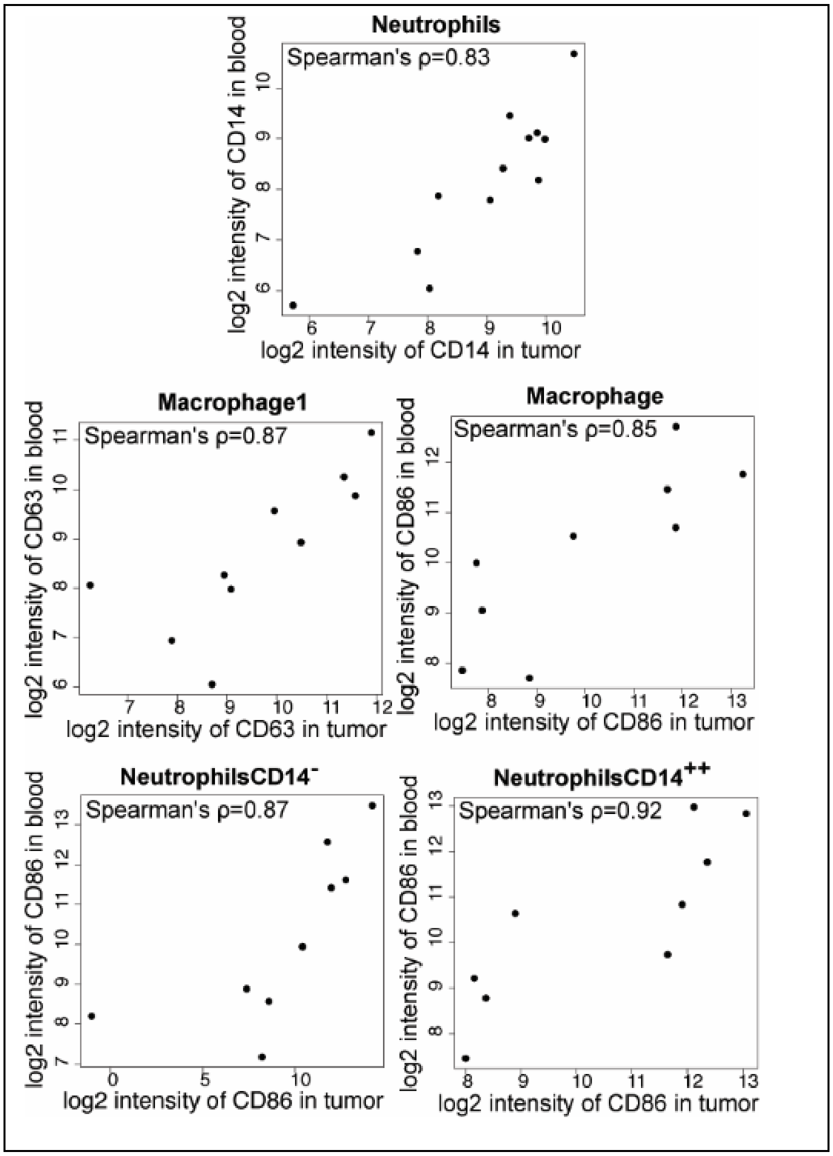


**Supplementary Figure 7:** Scatterplots for marker-cell type pairs which showed significant correlation between blood and tumor levels (supporting figure 5B)

### Supplementary Tables

**Supplementary Table 1:** Panel of antibodies used for staining of cells from blood and tumor tissue of individuals with glioma as well as healthy controls. Names of the surface proteins that were assessed along with clones (in parentheses) used are listed. Data for the names with highlighted text (in blue) is not presented as optimization of antibody concentration to be used did not result in positive binding (either the antibodies did not bind to their surface proteins in the protocol used (likely), or the cells did not express these markers under all conditions that were tested). All antibodies were purchased from BD biosciences.

| **Fluorophore** | **Panel 1** | **Panel 2** | **Panel 3** | **Panel 4** |
|---|---|---|---|---|
| Brilliant Violet 421 | CD294 | CD68 (Y1/82A) | CD69 (FN50) | CD33 (WM53) |
| Brilliant Violet 510 | Live-Dead | | | |
| Brilliant Violet 605 | CD14 (M5E2) | | | |
| Brilliant Violet 650 | CD15 (HI98) | | | |
| Brilliant Violet 786 | CD62L (SK11) | CD284 (TF901) | CD274 (M1H1) | CD80 (L307.4) |
| FITC / Alexa Fluor 488 | CD16 (3G8) | CD282 (11G7) | CD54 (HA58) | CD86 (2331-FUN-1) |
| PE | CD63 (H5C6) | CD36 (CB38) | VEGFR2 (89106) | CD163 (GHI/61) |
| PE-CF594 | - | HLA-DR (G46-6) | - | HLA-DR (G46-6) |
| APC / Alexa Fluor 647 | CD66b (G10F5) | - | - | CD206 (19.2) |
| Alexa Fluor 700 | - | CD11b (ICRF44) | - | CD11b (ICRF44) |
| APC-Cy7 | CD45 (2D1) | | | |

**Supplementary Table 2:** Clinical and sample collection details of data reported in this study. IHC =immunohistochemistry

| ID No. | Histology based grading | IDH status | Steroidal drugs administered prior to surgery | IHC performed | Availability for Flow Cytometry | | Other health issues |
|---|---|---|---|---|---|---|---|
| | | | | | Blood | Tumor | |
| MN-317 | GBM | negative | Yes | Yes | Yes | Yes | Diabetes |
| MN-321 | GBM | negative | Yes | Yes | Yes | Yes | |
| MN-322 | Grade III | positive | Yes | Yes | Yes | Yes | |
| MN-327 | GBM | negative | Yes | Yes | Yes | Yes | |
| MN-357 | GBM | negative | Yes | Yes | Yes | Yes | Recurrent GBM. Diagnosed with pericardial effusion, sleep apnea, and chronic papilloedema. Individual was on Wysolone for 2 years pre-surgery and special clearance was received for surgery |
| MN-363 | Grade III | positive | Yes | Yes | Yes | Yes | |
| MN-372 | Grade III | positive (weakly) | Yes | Yes | - | Yes | |
| MN-392 | GBM | negative | Yes | Yes | Yes | Yes | |
| MN-457 | Benign | - | Yes | - | Yes | Yes | |
| MN-458 | Grade III | positive | Yes | - | Yes | Yes | |
| MN-460 | GBM | negative | Yes | - | Yes | - | |
| MN-461 | Grade III | positive | Yes | Yes | Yes | Yes | |
| MN-474 | GBM | negative | Yes | - | Yes | Yes | Hypertension (diagnosed at least 8 years prior to surgery) |

| | | | | | | | |
|---|---|---|---|---|---|---|---|
| MN-478 | GBM | negative | Yes | - | Yes | Yes | On treatment for Diabetes Mellitus and Hypertension. Glaucoma surgery 6 months prior to tumor resection surgery |
| MN-494 | Grade III | positive | Yes | Yes | Yes | Yes | |

**Supplementary Table 3:** Subsets of cell types obtained from single-cell analysis of flow cytometry data

| **Cell type** | **Label** |
|---|---|
| SSCA$^{++}$CD14$^{-}$CD15$^{+}$ | Granulocytes |
| SSCA$^{+}$CD14$^{-}$CD15$^{+}$ | Granulocytes |
| SSCA$^{-}$CD14$^{-}$CD15$^{++}$ | Immature neutrophils |
| SSCA$^{++}$CD14$^{++}$CD15$^{-}$ | Macrophage |
| SSCA$^{++}$CD14$^{++}$CD15$^{+}$ | Macrophage1 |
| SSCA$^{+}$CD14$^{++}$CD15$^{+}$ | Monocyte |
| SSCA$^{+}$CD14$^{++}$CD15$^{-}$ | Monocyte |
| SSCA$^{++}$CD14$^{+}$CD15$^{++}$ | Neutrophils |
| SSCA$^{+}$CD14$^{+}$CD15$^{++}$ | Neutrophils |
| SSCA$^{++}$CD14$^{-}$CD15$^{++}$ | NeutrophilsCD14$^{-}$ |
| SSCA$^{+}$CD14$^{-}$CD15$^{++}$ | NeutrophilsCD14$^{-}$ |
| SSCA$^{++}$CD14$^{++}$CD15$^{++}$ | NeutrophilsCD14$^{++}$ |
| SSCA$^{+}$CD14$^{++}$CD15$^{++}$ | NeutrophilsCD14$^{++}$ |
| SSCA$^{-}$CD14$^{-}$CD15$^{-}$ | SSCA$^{-}$CD14$^{-}$CD15$^{-}$ |
| SSCA$^{-}$CD14$^{-}$CD15$^{+}$ | SSCA$^{-}$CD14$^{-}$CD15$^{+}$ |
| SSCA$^{-}$CD14$^{+}$CD15$^{++}$ | SSCA$^{-}$CD14$^{+/++}$CD15$^{++}$ |
| SSCA$^{-}$CD14$^{++}$CD15$^{++}$ | SSCA$^{-}$CD14$^{+/++}$CD15$^{++}$ |
| SSCA$^{-}$CD14$^{++}$CD15$^{+}$ | SSCA$^{-}$CD14$^{++}$CD15$^{-/+}$ |
| SSCA$^{-}$CD14$^{++}$CD15$^{-}$ | SSCA$^{-}$CD14$^{++}$CD15$^{-/+}$ |
| SSCA$^{-}$CD14$^{+}$CD15$^{-}$ | SSCA$^{-}$CD14$^{+}$CD15$^{-}$ |
| SSCA$^{-}$CD14$^{+}$CD15$^{+}$ | SSCA$^{-}$CD14$^{+}$CD15$^{+}$ |
| SSCA$^{++}$CD14$^{-}$CD15$^{-}$ | SSCA$^{++}$CD14$^{-}$CD15$^{-}$ |
| SSCA$^{++}$CD14$^{+}$CD15$^{-}$ | SSCA$^{++}$CD14$^{+}$CD15$^{-}$ |
| SSCA$^{++}$CD14$^{+}$CD15$^{+}$ | SSCA$^{++}$CD14$^{+}$CD15$^{+}$ |
| SSCA$^{+}$CD14$^{-}$CD15$^{-}$ | SSCA$^{+}$CD14$^{-}$CD15$^{-}$ |
| SSCA$^{+}$CD14$^{+}$CD15$^{-}$ | SSCA$^{+}$CD14$^{+}$CD15$^{-}$ |
| SSCA$^{+}$CD14$^{+}$CD15$^{+}$ | SSCA$^{+}$CD14$^{+}$CD15$^{+}$ |

**Supplementary Table 4:** Correlation coefficients (Spearman's r) along with p value presented for comparison of MDSC percentages in blood and tumor.

| | **GBM** | | | **Grade III** | | |
|---|---|---|---|---|---|---|
| | **# pairs** | **Spearman r** | **p value** | **# pairs** | **Spearman r** | **p value** |
| MDSC (overall) | 4 | 0.80 | 0.33 | 3 | 0.0 | 1 |
| G-MDSC | 3 | 0.50 | 1 | 3 | 1 | 0.33 |
| M-MDSC | 3 | 1 | 0.33 | 3 | 1 | 0.33 |
